# Comparative of probiotics reveals cecal microbial component associated with performance parameters in broilers

**DOI:** 10.1101/846766

**Authors:** D.R. Rodrigues, W. Briggs, A. Duff, K. Chasser, R. Murugesan, C. Pender, S. Ramirez, L. Valenzuela, L.R. Bielke

## Abstract

Probiotics have become increasingly popular in poultry industry as a promising nutritional intervention to promote modulation of intestinal microbiota as a means of improving health and performance. This study aimed to determine the effects of different probiotic formulations on the cecal microbial communities and performance in 21 and 42 day-old-broilers, as well as to define associations between ceca microbial profile and growth parameters. Probiotics investigated included a synbiotic (SYNBIO), a yeast-based probiotic (YEAST), and three single-strain formulations of spore-forming *Bacillus amyloliquefaciens* (SINGLE1), *B. subtilis* (SINGLE2) and *B. licheniformis* (SINGLE3). Dietary inclusion of SYNBIO, YEAST, and SINGLE2 increased body weight (BW) by 7, 14, and 21d (*p*<0.05) compared to a basal diet without probiotics (CON). The treatments SYNBIO, and YEAST decreased mortality by 21d, while SYNBIO reduced the overall mortality rate by 42d (*p*<0.05). Bifidobacteriales had the highest (*p*<0.05) population in SINGLE2, whereas Clostridiales was reduced compared to CON, SINGLE1, and SINGLE3. The addition of SYNBIO into diet mainly stimulated (*p*<0.05) the cecal relative abundance of Lactobacillales by 21d. Besides, Spearman’s correlation analyses revealed that population of Lactobacillales was associated with lower Enterobacteriales, higher BW, and lower mortality of growing broilers. These results suggest that the modulation of ceca microbiota and the greatest productive parameters were achieved by supplementation of specific probiotic mixture. The selection of probiotics by their ability to drive cecal microbiota towards lactic acid bacteria colonization may be a strategic approach to improve the indicators of performance in broilers.

## Introduction

Worldwide, the decreased percentage of chickens treated with sub-therapeutic levels of antibiotics has attracted attention towards a better understanding of dietary alternatives as growth and health promoters. Among them, probiotics have been indicated as a promising nutritional intervention to manipulate the avian microbiome [1–4]. Beneficial bacteria colonization of intestinal microbiota is essential for favoring host growth and performance, while an unfavorable alteration of the commensal structure may promote enteric infections, thereby deteriorating welfare and the performance indicators of poultry production [5].

Probiotics have become increasingly popular across human medicine and livestock industry due to the following benefits in the host: stimulation of beneficial microbiota, reduction, and prevention of pathogen colonization, development of immune system, improvement in digestive efficiency, and maturation of intestinal microbiota [3,5–9]. Although several bacterial species and yeasts have been described as potential probiotic for broiler chickens; *Bacillus, Lactobacillus, Enterococcus, Bifidobacterium, Pediococcus*, and *Escherichia* are the most common bacterial genera used for probiotic formulations, whereas *Saccharomyces cerevisiae* is the most common yeast [5,7]. Some of the factors that have been claimed to be responsible for probiotic’s efficiency include the microbial viability in the gastrointestinal tract (GIT), the ability to adhere to epithelial cells and colonize the host GIT, capability to reproduce itself in the host, and production of metabolites [9,10].

However, there have been inconsistencies concerning the effectiveness of probiotic supplementation in shaping GIT microbial communities and promoting growth. Accordingly, a comprehension of how the microbiota profile modulated by probiotics affect the host phenotype is still needed. Therefore, the primary aim of this study was to determine the effects of different probiotic formulations on the cecal microbial communities and performance, as well as to define associations between ceca microbial profile and growth parameters of broiler chickens.

## Material and methods

### Experimental design and dietary treatments

A total of 720 one-day-old Ross 708 male chicks were allocated to 6 treatments in a completely randomized design. Eight replicates were assigned to each of the treatments with 15 birds per replicate. Treatments were based on supplemental diets including (1) basal diet without probiotics (CON); (2) Synbiotic (0.45 g/Kg; SYNBIO); (3) Yeast-based probiotic (1.12 g/Kg; YEAST); (4) Single-strain probiotic 1 (0.45 g/Kg; SINGLE1); (5) Single-strain probiotic 2 (0.27 g/Kg; SINGLE2) or (6) Single-strain probiotic 3 (0.45 g/Kg; SINGLE3).

The SYNBIO-based mixture was composed of 2 × 10^11^ CFU/g multi-species probiotic including *Lactobacillus reuteri, Enterococcus faecium, Bifidobacterium animalis, Pediococcus acidilactici*, and a prebiotic (fructooligosaccharide). The formulation YEAST was a non-bacterial probiotic-containing *Saccharomyces cerevisiae* (Moisture 11%, Crude fiber 25%). The single-strain probiotics were composed of spore-forming *Bacillus* spp. Formulation SINGLE1 contained 1.25 × 10^6^ CFU/g of *B. amyloliquefaciens*, while SINGLE2 comprised 10 billion spores/g of *B. subtilis*. Besides, each gram of the SINGLE3 contained 3.20 ×10^9^ CFU of *B. licheniformis*.

Birds were reared from 1 to 42d and housed in floor pens on fresh wood shavings litter with *ad libitum* access to a standard corn-soy diet and water [11]. The feeding program consisted of 3 phases: starter (1-7d), grower (8-21d), and finisher (22-42d). Stater diets were in mash form, whereas the grower and finisher diets were pelleted. All experimental procedures were approved by the Ohio State University’s Institutional Animal Care and Use Committee (IACUC).

### Growth performance

The birds were weighed individually weekly for the overall experimental period. Feed consumption for each pen was recorded by measuring feed residue on the same days as birds were weighed. Feed conversion ratio (FCR) was calculated as pen feed consumption divided by body weight gain per pen, corrected for mortality. Mortality was showed as cumulative mortality per treatment by 21 and 42 days of age.

### Sample collection and processing

To investigate the intestinal microbiota composition of probiotic-treated broilers on days 21 and 42, ceca were collected from four birds per pen for DNA extraction and next-generation sequencing (NGS). Cecal contents were weighed and mixed to create pooled samples from two birds (n=16 per treatment for each time collection) for DNA extraction. Next, 0.3g of the mixed digesta was added into a 2.0mL screwcap microcentrifuge tube with 0.2g of zirconia beads (0.1mm). DNA was extracted from each sample, along with pure culture bacterial samples, using the protocol from Arthur et al. [12] with several modifications. After extractions were completed, DNA quality and quantity were measured using a Synergy HTX, Multi-Mode Reader (BioTek, Winooski, VT), and all samples were diluted to a concentration of 20ng/µL for NGS analysis.

### 16S Sequencing Analysis

Generation of PCR amplicon was achieved by amplification of the V4-V5 region of the 16S rRNA gene using 515F and 806R primers (515F: GTGYCAGCMGCCGCGGTAA, 806R: GGACTACHVGGGTWTCTAAT). DNA samples were library prepared for NGS using the Illumina MiSeq platform (2 x 300 bp; Illumina, San Diego, CA, USA) by Ohio State University Molecular and Cellular Imaging Center.

A sequence quality screen was performed to ensure high-quality sequences were submitted to the analysis pipeline. Briefly stated, sequence quality was determined using the FASTQC and MultiQC toolkits. Sequence reads exhibiting a quality score of lower than 20 were removed. Further, low complexity reads, those shorter than 200 bp in length, and mismatched primers were also eliminated. Additionally, reads exhibiting low sequence qualities on either end were trimmed. The pre-processed FASTQ files were then imported to the QIIME2 platform for analysis. The main analytical steps were as follows: firstly, reads were de-multiplexed and classified into their respective samples. Next, additional sequence quality control measures and feature table construction were performed by the DADA2 algorithm implementation in QIIME2. Quality control measures eliminated reads with barcode errors, along with reads that had more than two nucleotides mismatches, and chimeras. The high-quality sequences originating from the afore-mentioned quality control measures were subsequently clustered together using the q2-feature-classifier plugin with the GreenGenes 13.8 reference database. The resulting feature table was used to calculate phylum and order-level abundance infographics.

### Statistical Analysis

All data were subjected to Analysis of Variance (ANOVA) as a completely randomized design using the JMP Pro13 Software (JMP Software, SAS Inc., 2018). Body weight (BW), Feed intake (FI) and FCR were compared using Student’s *t*-test (*p* ≤0.05) to determine differences across groups. The mean relative abundances of microbial communities were also compared with a Student’s *t*-test (*p* ≤0.05). For mortality, data were analyzed using the Chi-Square test (*p* ≤ 0.05) in SAS (SAS Inc., 2018). Additionally, the Spearman’s rank correlation coefficient (R) was applied to identify correlations between bacterial colonization patterns and performance parameters (R software version 3.4.1).

## Results

### Growth performance parameters

Dietary inclusion of SYNBIO, YEAST, and SINGLE2 increased BW by 7, 14, and 21d compared to CON (*p*<0.05; Fig 1).

**Fig 1.**
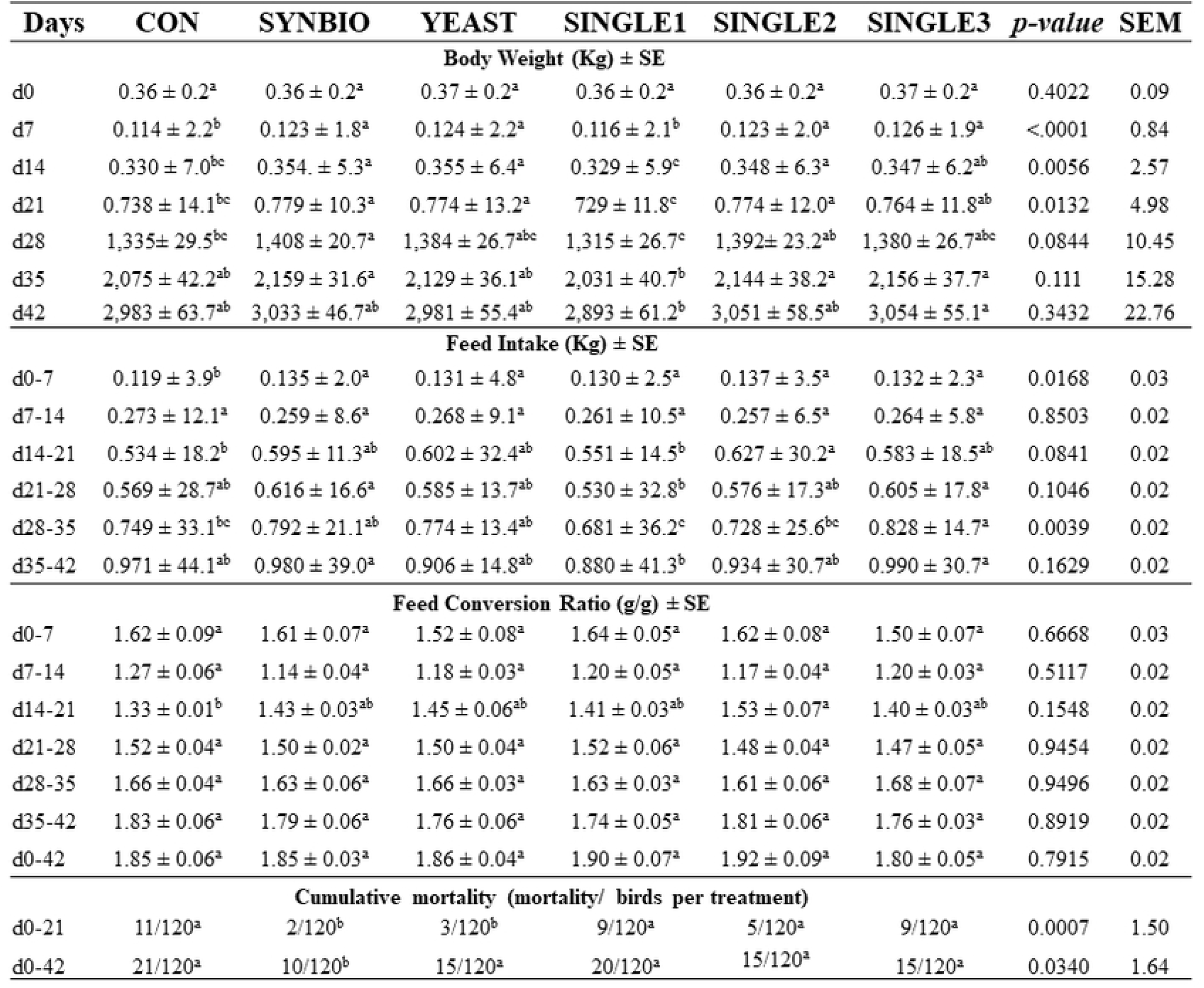
Performance parameters of broilers fed different probiotics from 1 to 42 days of age. ^a-c^ Different letters in the same row indicate statistical differences (*p*<0.05, Student’s t-test). Broilers fed basal diet without probiotics (CON), synbiotic (SYNBIO), yeast-based probiotic (YEAST), or single-strain formulations composed of *B. amyloliquefaciens* (SINGLE1), *B. subtilis* (SINGLE2), and *B. licheniformis* (SINGLE3).

In addition, broilers fed SYNBIO were heavier (*p*<0.05) than CON and SINGLE1 on day 28. By 35d and 42d, no significant differences were found in BW when compared against CON. Supplementation of probiotics in the diet increased (*p*<0.05) FI during d1-7 (Fig 1). However, there was no significant difference in FI between probiotic treatments and CON during all growth periods, except for d28-35, in which there was a significant increase of FI in SINGLE3. Similarly, no significant effect of probiotic supplementation was observed in FCR during d1-14 and d21-42. From d14 to 21, SINGLE2 had a statistically higher FCR (*p*<0.05) related to CON (Fig 1).

There was lower cumulative mortality in SYNBIO and YEAST treated birds by 21d. On 42d, dietary inclusion of SYNBIO significantly reduced the rate of mortality (*p*=0.03; Fig 1).

### Microbiota Composition

A total of 5,348,269 16S rRNA sequence reads were obtained. The number of mapped sequence reads of overall samples ranged from 13,545 to 60,125, with a mean of 27,855.82. In order to assess the impact of different probiotics supplementation on cecal bacterial populations, 16S-derived microbial community was analyzed at the taxonomic rankings of phylum and order levels.

Similar to many microbiome previous studies, the dominant cecal population mapped to the Firmicutes, Actinobacteria, and Proteobacteria (Table in S1 Table). On day 21, the relative abundance of the Actinobacteria phylum was statistically higher in SINGLE2 than CON and YEAST (Fig 2 A).

**Fig 2.**
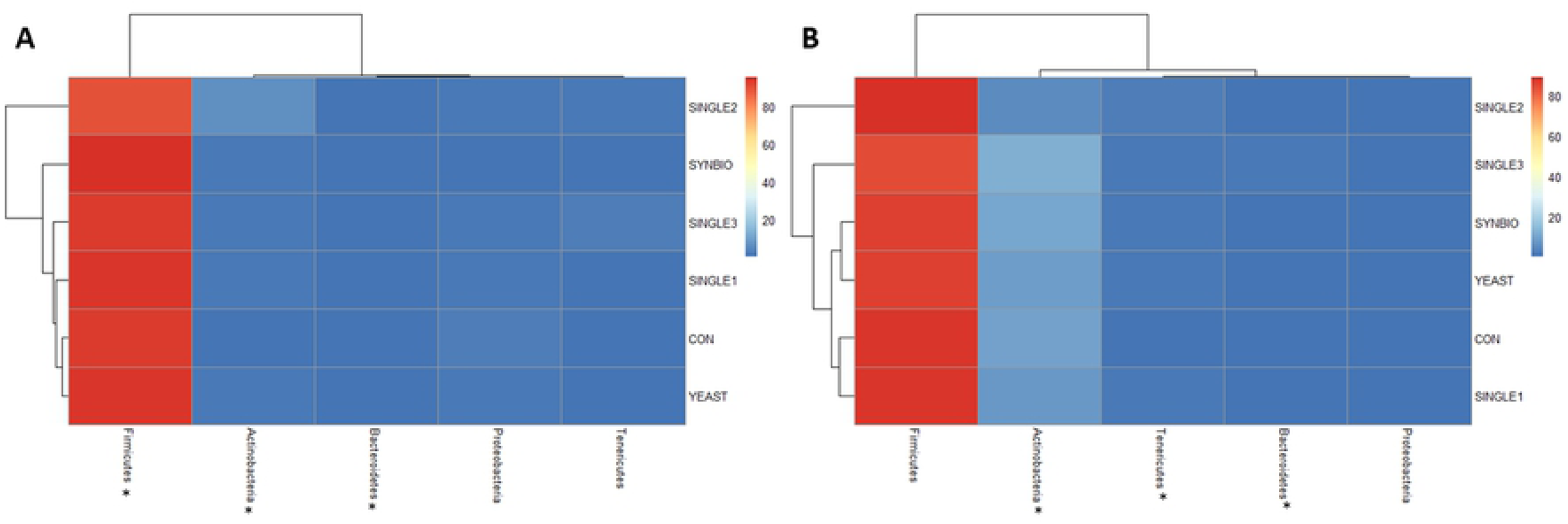
Cecal bacterial abundance at phylum-level of broilers fed different probiotics by 21 and 42 days of age. (A) Hierarchical clustering shown in a heat map of microbial communities profiles of samples from broilers fed basal diet without probiotics (CON), synbiotic (SYNBIO), yeast-based probiotic (YEAST), or single-strain formulations composed of *B. amyloliquefaciens* (SINGLE1), *B. subtilis* (SINGLE2), and *B. licheniformis* (SINGLE3) by 21 and (B) 42 days of age.. Statistical differences (*p*<0.05) between groups were reported for each bacterial population (*).

Similarly, Bifidobacteriales order had the highest (*p*<0.05) population in SINGLE2 (Fig 3A).

**Fig 3.**
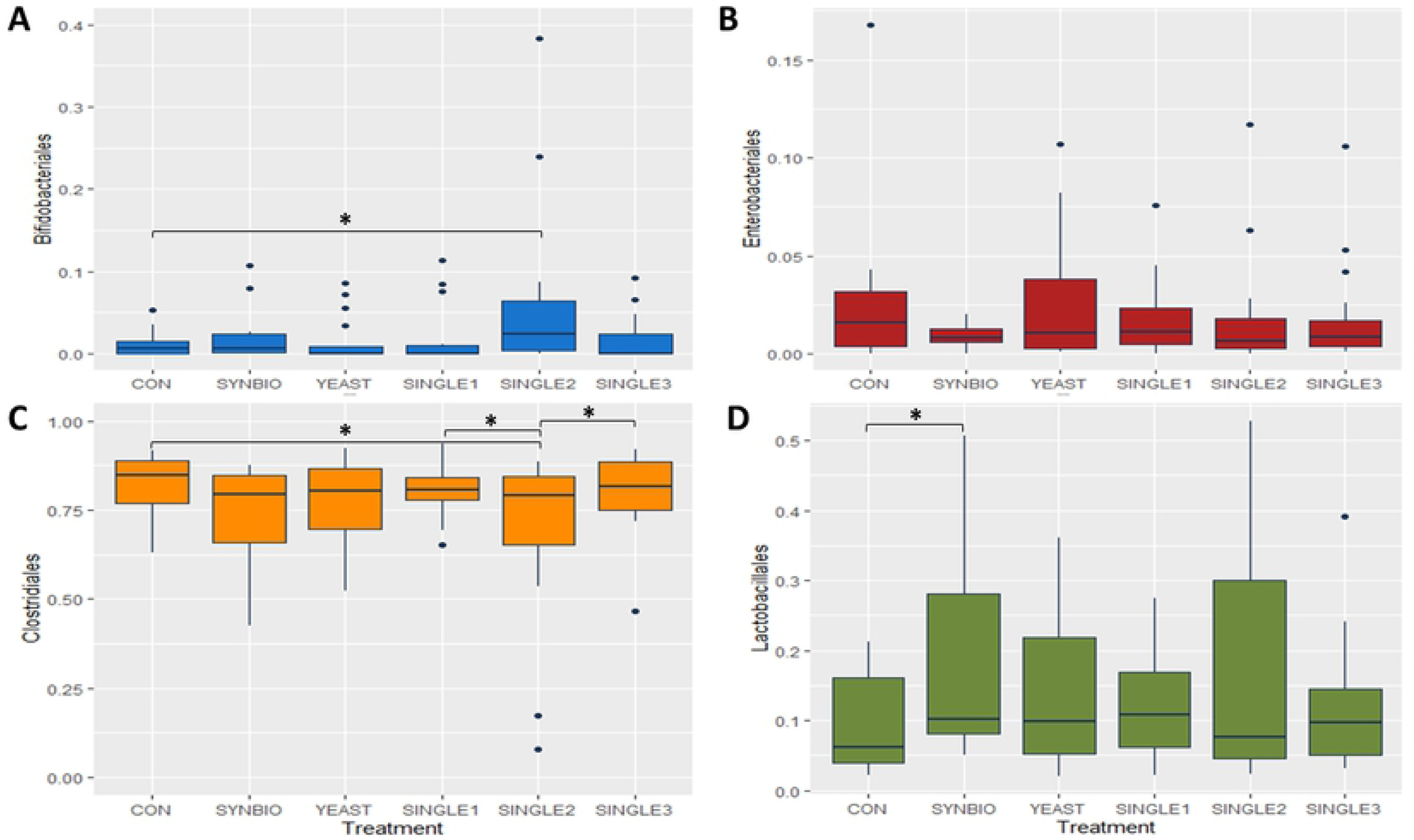
Microbial composition in the cecum digesta of 21-day-old broilers. Box plots show the relative abundance of the top four order-level bacterial population found in the ceca of broilers, including (A) Bifidobacteriales, (B) Enterobacteriales, (C) Clostridiales, and (D) Lactobacillales. The (*) at the top of the index represents significant differences between treatments (*p*<0.05).

There were significant changes concerning the Firmicutes phylum population at d21, which was reduced in SINGLE2 compared to CON and other treatments (Fig 2A). Clostridiales population was lower in ceca of SINGLE2 than CON, SINGLE1, and SINGLE3 (Fig 3C). In addition, supplementation of SYNBIO mainly stimulated (*p*<0.05) the cecal relative abundance of Lactobacillales order and decreased the Bacteroidetes phylum abundance compared to CON (Fig 3D; Table in S1 Table).

Bacteria belonging to the Proteobacteria phylum and Enterobacteriales order were not affected (*p*>0.05) by probiotic supplementation (Fig 2A), although a decrease in the relative abundance of these microbial communities were observed at 42d (Fig 2B).

Dietary treatments had minimal effects on cecal microbiota by 42 days of age. It may be noted that the Tenericutes population was increased in YEAST, SINGLE1, SINGLE2, and SINGLE3 (*p*<0.05). Besides, SINGLE2 had a lower abundance of Actinobacteria than SINGLE3, but no differences compared to CON were observed (Fig 2B)

### Correlation between microbiota composition and performance parameters

To further analyze the associations between cecal microbiome and host performance parameters, we conducted Spearman’s correlation linking the four discrepant order-level microbial taxa, BW, and mortality by 21 and 42 days of age. The Spearman’s rank correlation showed that abundances of Lactobacillales were negatively associated with Clostridiales by 21 days of age (Fig 5A).

**Fig 4.**
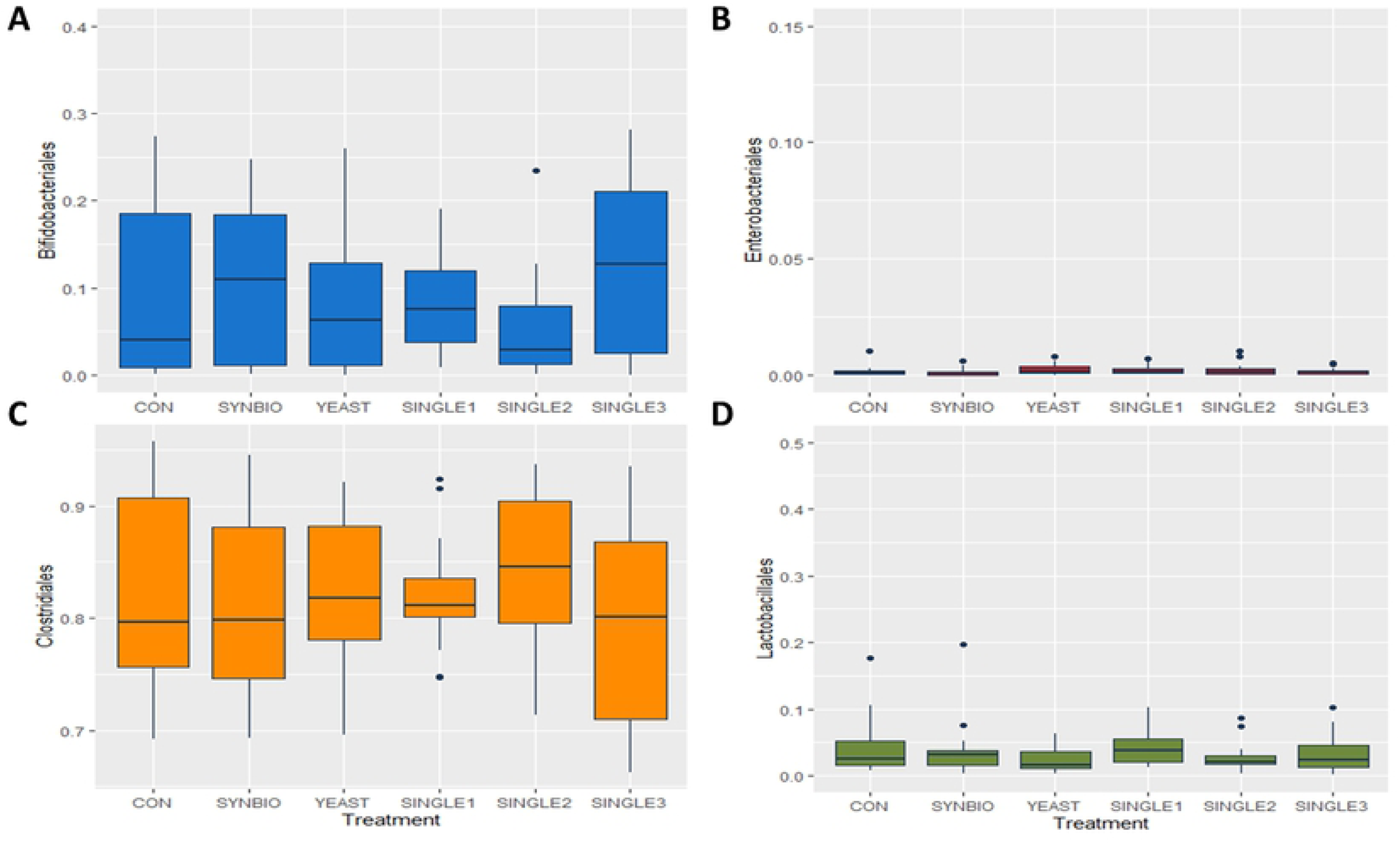
Order-level taxonomic distribution among samples from cecal contents of 42-day-old broilers. Bars represent the mean relative percentage of each bacterial population within samples from broilers treated with a basal diet without probiotics (CON), synbiotic (SYNBIO), yeast-based probiotic (YEAST), or single-strain formulations composed of *B. amyloliquefaciens* (SINGLE1), *B. subtilis* (SINGLE2), and *B. licheniformis* (SINGLE3). (A) Bifidobacteriales, (B) Enterobacteriales, (C) Clostridiales, and (D) Lactobacillales. The (*) at the top of the index represents significant differences between treatments (*p*<0.05).

**Fig 5.**
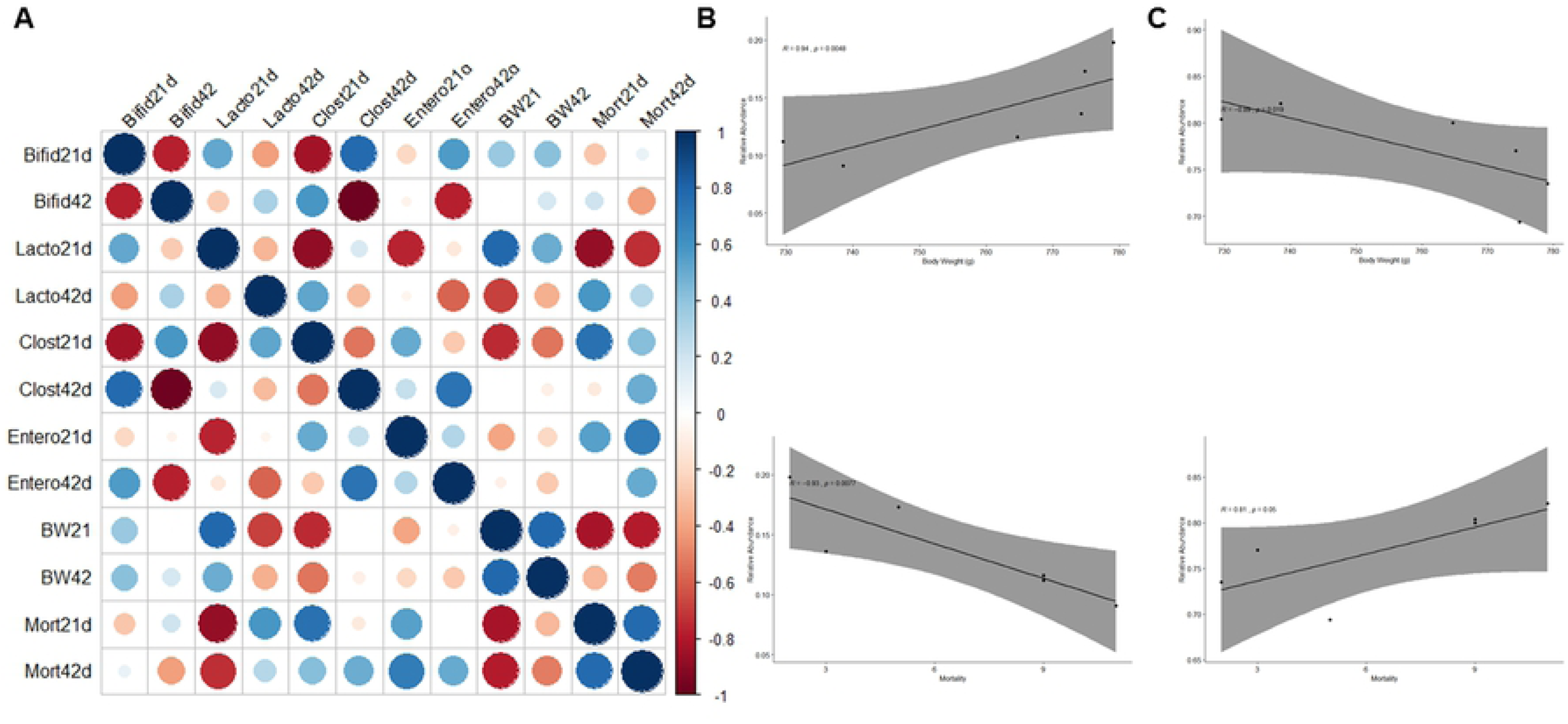
Spearman’s rank correlation matrix of the dominant microbial populations and growth performance parameters. (A) Strong correlations are indicated by large circles. The colors of the scale bar denote the nature of the correlation with 1 indicating a perfect positive correlation (dark blue) and −1 indicating perfect negative correlation (dark red). (**B**) Lactobacillales population by 21d was positively correlated with (R=0.94, *p*=0.048) Body Weight (BW) and negatively (R=-0.93, *p*=0.007) related to mortality rates (Mort). (**C**) At the same age, Clostridiales was negatively (R=-0.89, *p*=0.01) related to BW and positively (r=0.81, *p*=0.05) associated with mortality. Only significant Spearman’s correlation coefficients were screened.

Another interesting interaction between components of microbiota was the negative correlation among Enterobacteriales and Lactobacillales. Likewise, according to these analyses, the BW at d21 positively impacted BW at d42, and negatively influenced the mortality rate at both ages. A strong positive correlation between the cecal Lactobacillales population and BW (R=0.94, p=0.048; Fig 5B) at 21d was observed. At the same age, Lactobacillales’ relative abundance was negatively associated with the mortality rate (R=-0.93, *p*=0.007). These results indicate that a higher population of Lactobacillales in the ceca may be a marker of better performance for young broilers (21 days of age).

Similar associations were found within the Clostridiales population, which was negatively related to BW (R=-0.89, *p*=0.01; Fig 5C) and positively associated with mortality by 21 days of age. Interestingly, the greatest correlation was only identified at an earlier age.

## Discussion

This study was conducted in order to gain a better comprehension of how different probiotic formulations could modulate the cecal microbiota and impact on the indicators of performance in broilers throughout 42 days of age. Besides, the objective of the present investigation was also to identify an association between GIT microbiota phenotype and growth parameters. Our findings suggest that the greatest productive parameters of 21-day-old broilers promoted by specific probiotic-based supplementation were associated with a cecal microbial component.

The addition of probiotics into the diets supported a significant stimulation of FI and BW by 7 days of age. Probiotics seemed to have the greatest effect during the initial development of the microbiota [13]. As a consequence of limited contact with the hens’ microbiota, the assembly of the intestinal microbiome of the newly hatched chicks is predominantly influenced by the hatchery and farm environment [14–17]. Thus, an immediate supplementation of probiotics post-hatch is more important in avian species than in other animals [7]. The early exposure to microbial preparations has been identified as an approach to modulate the microbiota towards beneficial bacterial growth [3,4] and pathogen colonization reduction [18]. Additionally, supplementation of probiotics has been successfully linked to GIT development by stimulating the growth of villus surface area [18,19]. Other probiotic action mechanisms include maturation of immune system, improvement of gut barrier function, and the presence of highly competitive microbial communities, which can lead exclusion of pathogenic bacteria through competitive exclusion [3,5–9].

By 21 days of age, the probiotic mixture played a key role in improving bird BW. Dietary inclusion of YEAST and the bacteria-based probiotic SYNBIO consistently outperformed the CON birds. Although SINGLE2 increased BW at the same age, the feed efficiency (FI and FCR) was deteriorated compared to CON-treated broilers. Dietary supplementation of live yeast, yeast cultures, or yeast cell wall products was shown to have positive effects on animal performance [20–23]. Similar to our findings, Yalçın et al., 2013 showed that *Saccharomyces cerevisiae* supplementation improved weight gain during the starter period of broiler chickens, although there were no effects on final weight (42d). The performance improvements seen in SYNBIO were previously reported by Eckert et al., 2010. Enhanced performance promoted by synbiotic products may be related to the improvement in nutrient absorption, reduction of pathogens colonization, and stimulation of the immune response [19,24,25].

The results found in this study demonstrated that there was a treatment-specific effect on microbiota profiles, particularly evident in birds fed SINGLE2 and SYNBIO probiotics. It has been thought that many factors, such as the early intestinal colonization, physiologic stage of chickens, diet, or environment, can drive the composition and diversity of intestinal microbial communities [2–4,15,26,27]. The findings achieved here have shown that particular probiotic mixtures may also have benefits in modulating the microbiota of broilers. Of relevance, the supplementation of SYMBIO resulted in a robust modulation of intestinal microbiota with a high population of Lactobacillales, which may be explained by the ability of the lactic acid bacteria (LAB) strains, supplemented in the feed, to colonize and persist in the GIT. Moreover, the addition of prebiotics into the mixture may support the growth and activity of the probiotic and GIT beneficial bacteria [28]. Similar results were found in layers, in which the addition of SYNBIO in the feed increased the relative abundance of LAB in ceca showing that the supplemented strains survived and colonized the GIT [19]. Nevertheless, probiotics can also affect the development of the microbiota without effectively colonizing it by merely passing through the intestinal tract [15]. In this context, although the Bacillales did not become a resident of the cecal microbiome, it is possible that the supplementation of SINGLE2 possibly created a favorable environment for the Bifidobacteriales grow.

We further looked for associations between the cecal predominant microbial signature and indicators of growth parameters. Lactobacillales population in ceca was positively correlated to BW at 21d and negatively associated with mortality rate. The association between improved weight gain and LAB has also been observed by Yan et al., 2017 and De Cesare et al., 2019. Hence, LAB may be able to enhance the energy and mineral recovery from nutrients; its higher intestinal colonization results in a better digestive efficiency [29,31]. It is likely that the high abundance of Lactobacillales found in SYNBIO may have contributed to increasing the BW at 21d and 28d, as well as decreasing the overall number of dead birds in this treatment.

Spearman’s correlation analyses also revealed that the Clostridiales was negatively related to BW by 21d. This association was highly evident in SINGLE2 treated birds, which had the lowest Clostridiales population and increased BW compared to CON group. Clostridiales were the dominant order accounting for almost 83% of the entire cecal microbiota among treatments. Although Clostridiales members are known as the main responsible for short-chain fatty acid metabolism in chicken cecum (Oakley et al., 2014; Pandit et al., 2018), obtaining insights into how higher diversity and high colonization of other bacterial communities in ceca can influence growth parameters may have important implications for selecting probiotic formulations.

The results presented here are evidence that supplementation of specific probiotics mixtures, particularly seen with SYNBIO, can modulate the microbiota that colonizes the gut shortly after hatch, thereby influencing the performance and survival of chicks during their growth. Unlike the grower phase, the supplementation of probiotics did not affect the performance or cecal profile by 42d. It is worth highlighting that these chickens were under experimental conditions without a pathogen challenge or stress induction. It would be interesting for future studies to evaluate the tested probiotic formulations under challenge conditions to determine if the ongoing administration is necessary to affect microbiota profile and performance.

In conclusion, this study illustrated that not all probiotic-based formulations modulated the ceca microbiota to a similar extent, nor resulted in improved performance. The population of Lactobacillales was identified to be strongly associated with lower Enterobacteriales, higher BW, and lower mortality of growing broilers. Accordingly, the selection of probiotic mixture by their ability to drive cecal microbiota towards LAB colonization may be a strategic approach to improve the indicators of performance in broiler chickens.

## Supporting information

**S1 Table. Phylum-level taxonomic of cecal microbial communities in broilers at 21 and 42 days of age**

**S2 Table. Spearman’s rank correlation matrix of the dominant microbial populations and growth performance parameters**

